# Antagonistic plants preferentially target arbuscular mycorrhizal fungi that are highly connected to mutualistic plants

**DOI:** 10.1101/867259

**Authors:** Sofia IF Gomes, Miguel A Fortuna, Jordi Bascompte, Vincent SFT Merckx

## Abstract

- How antagonists – mycoheterotrophic plants that obtain carbon and soil nutrients from fungi – are integrated in the usually mutualistic arbuscular mycorrhizal networks is unknown. Here, we compare mutualistic and antagonistic plant associations with arbuscular mycorrhizal fungi and use network analysis to investigate fungal association preferences in the tripartite network.
- We sequenced root tips from mutualistic and antagonistic plants in a tropical forest to assemble the combined tripartite network between mutualistic plants, mycorrhizal fungi, and antagonistic plants. We compared the fungal ecological similarity between mutualistic and antagonist networks, and searched for modules (an antagonistic and a mutualistic plant interacting with the same pair of fungi) to investigate whether pairs of fungi simultaneously linked to plant species from each interaction type were overrepresented throughout the network.
- Antagonistic plants interacted with approximately half the fungi detected in mutualistic plants. Antagonists were indirectly linked to any of the detected mutualistic plants, and fungal pairwise ecological distances were correlated in both network types. Moreover, pairs of fungi sharing the same antagonistic and mutualistic plant species occurred more often than expected by chance.
- We hypothesize that the maintenance of antagonistic interactions is maximized by targeting well-linked mutualistic fungi, thereby minimizing the risk of carbon supply shortages.

## Introduction

Since the early history of life, interspecific mutualisms have been paramount in the functioning of ecosystems (Thompson, 2005; Bascompte & Jordano, 2013). Mutualisms can form complex networks of interdependence between dozens or even hundreds of species. A prime example of this ‘web of life’ is the 450-million-year-old mutualism between the majority of land plants and arbuscular mycorrhizal (AM) fungi (Strullu-Derrien *et al*., 2018). In this interaction plants supply the arbuscular mycorrhizal Glomeromycotina and Mucoromycotina fungi with carbohydrates, essential for fungal survival and growth. In return, the fungi provide their host plants with mineral nutrients and water from the soil (Smith & Read, 2008; Bidartondo *et al*., 2011). One of the key characteristics of the arbuscular mycorrhizal interaction is its low interaction specificity: a mycorrhizal plant typically associates simultaneously with multiple fungi and a mycorrhizal fungus often associates simultaneously with multiple plants (Lee *et al*., 2013). This creates complex underground networks in which plants of different species are indirectly linked through shared arbuscular mycorrhizal fungi (Toju *et al*., 2015; Chen *et al*., 2017). Despite this low specificity, there is evidence that networks of plants and arbuscular mycorrhizal fungi do not assemble randomly, and effects of plant functional group (Davison et al. 2011; Sepp *et al*., 2019), and plant or fungal evolutionary relationships (Montesinos-Navarro *et al*., 2015; Chen *et al*., 2017) on arbuscular mycorrhizal interaction networks have been reported.

Considerable progress has been made in dissecting the exchange of resources between plants and AM fungi, and its regulation, in recent years. Experimental work on low-diversity systems has demonstrated that control is bidirectional, and partners offering the best rate of exchange are rewarded, suggesting that AM networks consist of ‘fair trade’ interactions (Bever *et al*., 2009; Kiers *et al*., 2011). However, the importance of reciprocally regulated resource exchange is questioned, as mycorrhizas also affect plant health, interactions with other soil organisms, host-defense reactions, and suppression of non-mycorrhizal competitor plants (Walder & Van Der Heijden, 2015). Also, strictly reciprocal regulation of resource exchange does not seem to apply to all AM interactions. Some exceptional plants, for example, behave as antagonists (Selosse & Rousset, 2011; Walder & Van Der Heijden, 2015): mycoheterotrophic plants can obtain carbon from root-associated fungi and some species have replaced photosynthesis by carbon uptake from arbuscular mycorrhizal fungi (Leake, 1994; Merckx, 2013). The mechanism underpinning carbon transfer from arbuscular mycorrhizal fungi to mycoheterotrophic plants remains unclear, but it is unlikely that resource exchange in these mycoheterotrophic associations relies on reciprocal dynamics. Hence, mycoheterotrophic plants are considered cheaters of the mycorrhizal symbiosis because they exploit the arbuscular mycorrhizal symbiosis for soil nutrients and carbon without reciprocating, to our current knowledge (Selosse & Rousset, 2011), or without apparently being sanctioned by the fungal partners (Walder & Van Der Heijden, 2015). Moreover, it has been suggested that mycoheterotrophic plants may display a truly biotrophic parasitic mode, digesting the fungus colonizing their roots (Imhof *et al*., 2013).

Phylogenetic studies demonstrate that these cheaters evolved from mutualistic ancestors (Merckx *et al*., 2013). Within obligate mutualisms, the critical barrier to mutualism breakdown and to the evolutionary stability of the resulting cheater species is thought to be a requirement for three-species coexistence: a cheater plant relies on a mutualistic partner – a mycorrhizal fungus – which simultaneously interacts with a mutualistic plant (Pellmyr & Leebens-Mack, 1999). In species-rich mutualisms such as the arbuscular mycorrhizal symbiosis, where multi-species coexistence is the rule, a high potential for the occurrence of these tripartite linkages is expected (Merckx & Bidartondo, 2008). Indeed, while evolution of cheating in specialized obligate mutualisms is relatively rare (Sachs & Simms, 2006), cheating in the arbuscular mycorrhizal symbiosis evolved in more than a dozen of plant clades, including over 250 species which together occur in nearly all tropical and subtropical forests (Merckx, 2013; Gomes *et al*., 2019a).

Previous work has shown that mycoheterotrophic plants – hereafter ‘antagonists’ – target a subset of the mycorrhizal fungi available in the local community (e.g Bidartondo *et al*. (2002); Gomes *et al*. (2017a); Sheldrake *et al*. (2017)). Their associated fungal communities can vary in specificity (Merckx *et al*., 2012), and in communities of co-occurring antagonists the phylogenetic diversity of their fungal associations appears to increase proportionally to their fungal overlap, a pattern which potentially responds to an ecological mechanism driven by maximizing co-occurrence and avoiding competitive exclusion among antagonistic plants. (Gomes *et al*., 2017b). However, whether partner choice of these antagonists is affected by mutualistic mycorrhizal interactions is currently unknown. Here, we hypothesize that antagonists preferentially associate with ‘keystone’ (Mills & Doak, 1993) fungi that are connected to many different mutualistic plants, since these fungi are potentially more resilient to perturbations (Bascompte & Jordano, 2007) and may be the most reliable source of carbon (Waterman *et al*., 2013). In addition, since fungal traits play an important role in arbuscular mycorrhizal interactions – phylogenetically related AM fungi (assumed to have similar functional traits), preferentially interact with similar plant species (Chagnon *et al*., 2015) – we hypothesize that if partner selection in tripartite networks is trait-driven we will be able to detect an influence of the phylogenetic relationships of the fungi. Here, we test these hypotheses on a combined tripartite mycorrhizal network of co-occurring antagonistic and surrounding mutualistic plants linked by shared arbuscular mycorrhizal fungi compiled by high-throughput DNA sequencing.

## Materials and methods

### Sampling

Since mycoheterotrophic plants are relatively rare and often have patchy distributions (Gomes *et al*., 2019b), we sampled two 4 x 4 m subplots a few meters apart in a coastal low land plain rainforest in French Guiana (5°28’25”N 53°34’51”W) on 28 July 2014, with overlapping antagonistic species. In both subplots, roots of antagonistic plants and surrounding mutualistic plants were sampled, cleaned with water, and stored on CTAB buffer at -20oC until further processing. We found in total five antagonistic plant species: *Dictyostega orobanchoides, Gymnosiphon breviflorus* (Burmanniaceae), *Voyria aphylla, Voyriella parviflora* (Gentianaceae), and *Soridium spruceanum* (Triuridaceae). Complete individuals of antagonistic plants were dug out, and around each, 3 root tips of mutualistic plants were collected. In addition, we sampled five additional mutualistic plant root tips randomly from each quadrant of each plot aiming to better represent the local belowground mutualistic community. A detailed taxon list of antagonistic and mutualistic plant species collected can be found in Supporting Information, Table **S1**.

### Plant identification and fungal communities sequencing

DNA was extracted from the CTAB-preserved roots with the NucleoMag96 Plant Kit (Macherey-Nagel Gmbh & Co., Düren, Germany), using the KingFisher Flex Magnetic Particle Processor (Thermo Fisher Scientific, Waltham, MA, USA). Mutualistic plant species were identified by sequencing the markers *matK or trnL* as described in (Gomes *et al*., 2017a). Plant identification to species, genus, or family level based on BLAST against GenBank was reviewed by consulting the checklist of plants in French Guiana (Berry & Weitzman, 2007). Fungal communities associated with the roots of mutualistic and antagonistic plants were amplified using the primers fITS7 (Ihrmark *et al*., 2012) and ITS4 (White *et al*., 1990) and sequenced with a Personal Genome Machine (Ion Torrent; Life Technologies, Guildford, CT, USA), as described in Gomes *et al*. (2017b) in two separate runs. Negative controls from the extraction and the PCR reactions were included (and had zero reads). The same bioinformatics methods from Gomes et al. (2017b) were used to process the raw reads from the two runs combined until clustering into 97% Operational Taxonomic Units (OTUs), using USearch v.7.0 (Edgar, 2010). OTUs represented by less than six reads in each sample were excluded to avoid spurious OTUs (Lindahl *et al*., 2013). The taxonomical assignment of each OTU was done by querying against the UNITE database (Kõljalg *et al*., 2013). Because the antagonistic plants in this study are presently only known to associate with fungi that belong to the sub-phylum Glomeromycotina (Merckx *et al*., 2012), we only retained fungal OTUs from this sub-phylum in the subsequent analysis. In these analyses, we accounted for the phylogenetic relatedness between fungal OTUs, due to the uncertainty of correspondence of individual OTUs with taxon diversity of arbuscular mycorrhizal fungi (Flynn *et al*., 2015), despite the fact that delimitation of OTUs is not likely to interfere with ecological interpretations (Lekberg *et al*., 2014). We inferred the phylogenetic relationships between the fungal OTUs following the strategy of Gomes *et al*. (2017b). Briefly, we aligned the OTUs with partial reference sequences and performed a phylogenetic inference in which the relationships between these references were enforced based on Krüger *et al*., (2012). Hereafter, we refer to OTUs as ‘fungi’. The fungal communities obtained from the root samples are not necessarily similar representations of the total communities of antagonists and mutualists because the sampling coverage of their root systems varied: antagonistic plants are small herbs, and thus large parts of their root system were collected and extracted, while for mutualistic plants, mostly forest trees (see Results), root samples represents only a small fragment of their root system and thus the detected fungal communities associated with mutualistic plants are likely a subset of their total fungal communities.

### Plant-fungal interactions

Fungal communities at the plant individual level were significantly structured by plant species identity with a minor influence of the subplot where they were collected (see Supporting Information, Methods **S1**, Figure **S1**). To contrast the interactions of antagonistic and mutualistic plants, we considered their plant-fungal interactions as a single tripartite network (Fig **1a**) by combining the fungal communities associated with individual plants from the two subplots into overall communities per plant species. By combining the data from the two subplots, we aimed to reconstruct a more robust picture of the interactions in this highly diverse rainforest while compensating for the relatively low sampling intensity (for 77 out of 220 root tips of mutualistic plants both plant identification and AM fungal community data with sufficient reads could be obtained). Plant species for which Glomeromycotina fungi were represented by less than 500 reads were excluded, resulting in removing 11 mutualistic plant species (see details in Supporting Information, Table **S1**). Moreover, because rarefying the OTU matrix has been shown to greatly increase the false positive rate of OTUs per sample (McMurdie & Holmes, 2014), we performed the subsequent analyses on 100 matrices rarefied to 844 reads per plant species, based on lowest number of reads obtained for all plant species in the observed dataset. Results are presented with mean and standard deviation values obtained from running the analyses on the rarefied matrices. To investigate fungal association patterns of antagonistic and mutualistic plants, we used incidence (binary) data, because our main interest was to determine which interactions can be established and not how abundant they are. Also, due to the difference in root sampling (see above) DNA read abundance data unlikely correspond to fungal abundances at the plant species level, in particular between plant types (antagonist and mutualists). We considered the simultaneous presence of a fungus in the roots of an antagonistic and a mutualistic plant as a potential link between both (Southworth *et al*., 2005).

**Figure 1.**
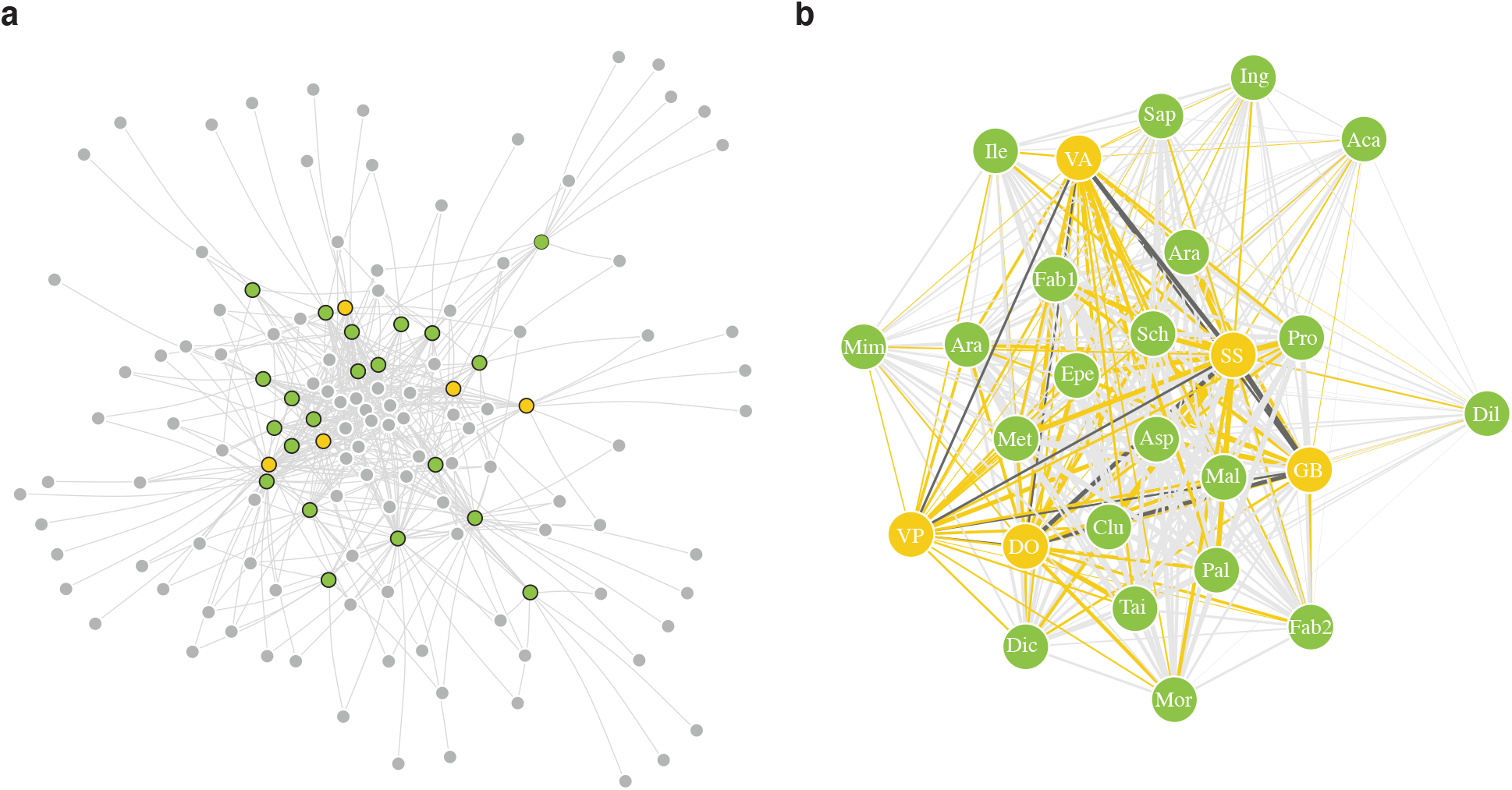
Tripartite arbuscular mycorrhizal interactions. (a) Visualization of the tripartite network between fungi (grey) and antagonist (yellow), and mutualist (green) plants, where edges represent a connection between a plant and a fungus. (b) Plant-plant fungal overlap network, in which edges represent a link between plant species through shared arbuscular mycorrhizal fungus. The thickness of the lines represents the interaction strength between the plants (the thicker, the more fungi are shared). Yellow lines link antagonistic and mutualistic plants; light grey lines link mutualistic to mutualistic plants; and dark grey lines link antagonistic to antagonistic plants. Identification of autotrophic plants is indicated by the first three letters of their name (see full names in Fig. 3); mycoheterotrophic plants are *S. spruceanum* (SS), *G. breviflorus* (GB), *D. orobanchoides* (DO), *V. aphylla* (VA), and *V. parviflora* (VP). In both network representations (a, b), one of the 100 rarefied matrices to a depth of 844 reads was used; and the Fruchterman-Reingold layout was used, in which nodes are evenly distributed through the graph, where plants that share more connections are closer to each other.

We tested for the effect of plant and fungal phylogenetic relatedness on the observed interactions in both networks. We used the fungal phylogeny described above, and the phylogenetic relationships of the plants as derived from *TimeTree* (Kumar *et al*., 2017) (see details in Supporting Information, Methods **S2**). We computed Mantel test correlations between the phylogenetic distance matrix and the community dissimilarity matrix in each instance for antagonistic and mutualistic species individually. The phylogenetic distance matrices were extracted from the phylogenetic trees of plants and fungi, and the community dissimilarity matrices were calculated as the Jaccard distance on the binary interaction matrices. Phylogenetic signal was calculated for the 100 rarefied matrices, and consistency of significant results was assessed across multiple rarefaction depths (Supporting Information, Fig. **S2**).

### Mutualistic and antagonistic plant-plant interactions

To compare the range of fungal interactions between mutualistic and antagonistic plants, we calculated their normalized degree and the phylogenetic species variability of their associated fungal communities. The normalized degree of a plant species is the proportion of its associated fungi out of the total possible fungi in the network (Martín González *et al*., 2010), and was calculated with the *ND* function of the *bipartite* R package (Dormann *et al*., 2018). The phylogenetic species variability (*psv*) of the fungal community associated with a plant species summarizes the level to which the fungi in this community are phylogenetically related (Helmus *et al*., 2007a). When a community consists of unrelated fungi, the index equals 1, indicating maximum phylogenetic variability. As relatedness increases, the index approaches 0, indicating high phylogenetic specificity (Helmus *et al*., 2007b). To explore how the number of interactions relates to the phylogenetic relationships among the detected fungi, we tested for a linear correlation between normalized degree and *psv* by calculating the Pearson’s correlation coefficient. In addition, we calculated the fungal overlap between each pair of plant species as the number of fungi they share to infer plant–plant preferences (Fig **1b**).

### Mutualistic and antagonistic fungal interactions

To investigate whether fungi have similar number of interactions with plants in both mutualistic and antagonistic networks, we calculated the Pearson correlation between the normalised degrees of the shared fungi within each interaction type. We compared the ecological similarity of the fungi shared between the two interaction type networks (i.e. mutualistic and antagonistic interactions). The ecological similarity of a pair of fungi represents their similarity in interactions through shared plants. We calculated similarity matrices between the fungi linked to mutualistic and antagonistic plants separately, using two different measures. The first measure was the Jaccard index, which corresponds to the number of plant species with which both fungi interact divided by the total number of plant species with which they interact, and the second was the Overlap measure: C_*ij*_/min(d_*i*_, d_*j*_), where C_*ij*_ is the number of shared plants between fungus *i* and *j*, and min(d_*i*_, d_*j*_) is the smallest number of associated plants between fungus *i* and *j* (Saavedra *et al*., 2013). The overlap measure corresponds to the number of shared plant species relative to the maximum number of plant species that can be shared, and takes into account the possibility that fungi have differential limits in their maximum number of plant partners. We computed the Mantel test correlation between the similarity matrices for the mutualistic and antagonistic interactions using the two different measures. We computed the partial Mantel test correlation between the similarity matrices calculated using the Jaccard and Overlap measures, controlling for the phylogenetic relatedness of the shared fungi.

### Network analysis

We searched for diamond-shape modules in the tripartite network (reduced to only include fungi shared between both interaction types) representing a mutualistic and an antagonistic plant interacting with the same pair of fungi (Fig. **2**). If the module is overrepresented throughout the entire network, it can be considered a motif (Milo *et al*., 2002; Bascompte & Melián, 2005; Stouffer *et al*., 2007). This specific motif does not consider the identity of the partners *per se*, but searches for this particular topological structure. To assess whether the diamond-shaped module was overrepresented relative to random expectations, we randomized the original antagonistic plants – mycorrhizal fungi – mutualistic plants community matrix and calculated the frequency of this diamond-shaped module in 1000 generated random networks. We used a null model that draws an interaction between a plant and a fungal species which is proportional to the generalization level of both species. Specifically, this probability is defined as the arithmetic mean of the fraction of interactions of the plant and that of the fungi (null model 2 in (Fortuna & Bascompte, 2006)), initially proposed in (Olesen *et al*., 2003). Because there was an imbalanced number of plant species (five antagonistic and 21 mutualistic plants), each randomized matrix resulted from randomizing the antagonistic and mutualistic interactions separately, and then combined into a single matrix before module search. To exclude the effects of sampling effort both in the number of individuals collected (leading to roughly half of fungal reads belonged to the five antagonistic species while the other half represented 21 mutualistic species) and in representation of whole (antagonists) or partial (autotrophic) root systems, we repeated this procedure in a set of 100 rarefied matrices where each set was resampled to multiple depths.

**Figure 2.**
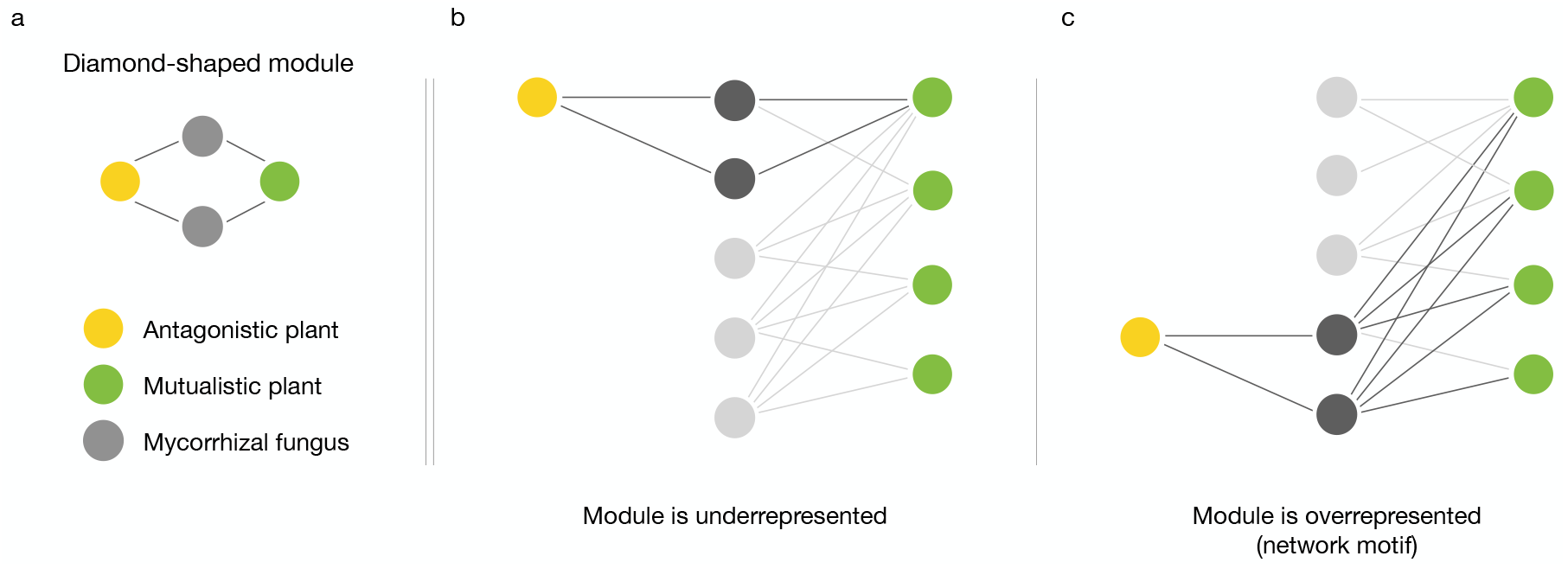
Diamond-shape module investigated in this paper. Representation of the module linking pairs of fungi to the same antagonistic and mutualistic plant species (a). Examples of under-(b) and overrepresentation (c) of the module in a network. When the module is overrepresented it is considered a network motif.

To evaluate whether the diamond-shaped module was overrepresented throughout the empirical network, we calculated the standard 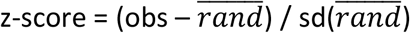, where *obs* is the observed network modules and 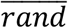 is the mean of the same type of modules across the 1000 random networks, using a 95% confidence interval. An empiric result above the null confidence interval indicates that pairs of fungi are shared between particular antagonistic and mutualistic plants more often than expected by chance, reflecting a preference of antagonistic plants to link to pairs of fungi that are generalists in their interactions with mutualistic plants. An empirical result below the null confidence interval indicates that antagonistic plants target pairs of fungi that are specialists in their interactions with mutualistic plants.

## Results

### Plant-fungal interactions

We successfully extracted DNA from root tips of 32 mutualistic (123 root tips) and five antagonistic (60 individuals) plant species (see Table **S1** for detailed species list). After retaining samples that contained a minimum of 500 reads identified as Glomeromycotina, we reduced our dataset to 21 mutualistic (77 root tips) and five antagonistic (27 individuals) plant species, with a total of 365,135 reads. Mutualistic plants belonged to 24 families, including 16 trees, two shrubs, one herb, one vine, and one liana. We obtained 115 arbuscular mycorrhizal fungi, identified as Glomeromycotina within three families Gigasporaceae (4 fungi), Acaulosporaceae (14) and Glomeraceae (97). Of these, 96 OTUs (49% of total reads) were present in the antagonistic network.

The 100 rarefied binary matrices included 26 plant species (21 mutualists and 5 antagonists) and 110 ± 1.47 fungi. Of these, mutualistic networks had 100 ± 1.51 fungi and antagonistic networks had 61 ± 3.14 fungi. In all the generated matrices, mutualistic and antagonistic plants shared 51 ± 3.16 fungi, which represents 55% and 92% of total fungi present in mutualistic and antagonistic plants, respectively (Fig. **1a**).

We measured a relatively low but significant phylogenetic signal of the fungal phylogeny on the antagonistic (*r* = 0.20 ± 0.06, *p* = 0.003 ± 0.005) and mutualistic (*r* = 0.21, *p* = 0.001) networks (see effect of rarefaction depth on phylogenetic signal in Supporting Information, Fig. **S2**). We found no significant effects of plant phylogeny on any of the networks.

### Mutualistic and antagonistic plant-plant interactions

The number of fungal interactions per plant species varied, with *Aspidosperma* sp. showing the highest normalised degree and Sapindaceae sp. the lowest amongst the mutualistic plants (Fig. **3a**). Ranked over all the plants, three antagonistic species, namely *V. aphylla, V. parviflora* and *D. orobanchoides*, presented a normalised degree in the lower half of the spectrum, while *G. breviflorus* and *S. spruceanum* are amongst the plants with the highest number of associated fungi. We also observed a wide range of phylogenetic species variability (psv) for mutualistic plants, with *Acacia* sp. and *Aspidorperma* sp. with the highest *psv* and Sapindaceae sp. the lowest, throughout which, the antagonistic plants are distributed (Fig. **3b**). There was a significant correlation between the normalised degree and *psv* of the mutualistic plants (Pearson’s correlation: *r* = 0.54, *t* = 2.80, *df* = 19, *p* = 0.011), while for the antagonistic plants the number of interactions (normalised degree) does not reflect the fungal diversity (*psv*) in their roots (Pearson’s correlation: *r* = -0.29, *t* = -0.53, *df* = 3, *p* = 0.632).

**Figure 3:**
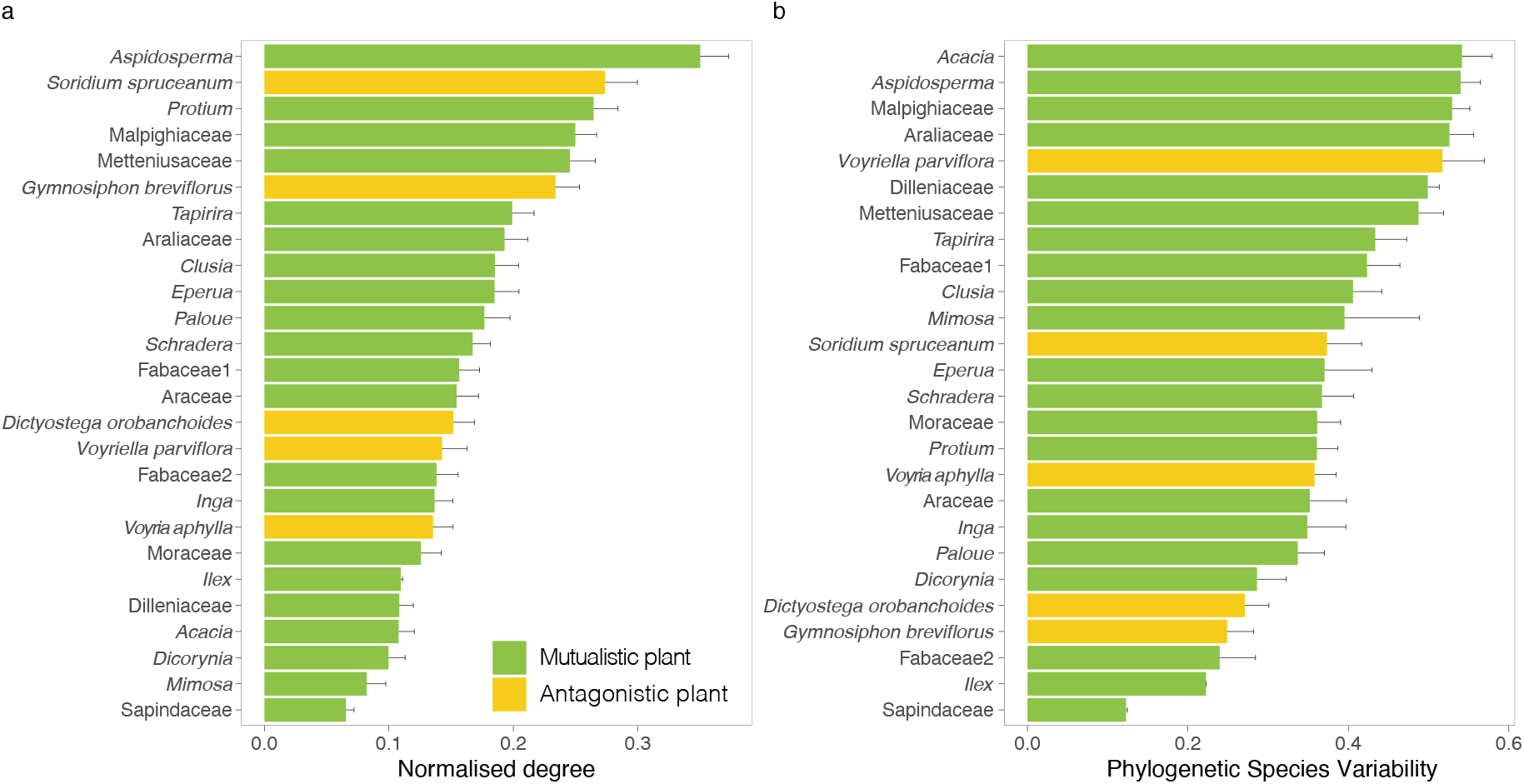
Mean ± SD plant normalised degree and PSV for the 21 autotrophic and five mycoheterotrophic plant species, resulted from the 100 rarefactions to a depth of 844 reads.

Within the antagonistic plants, we observed that *S. spruceanum* shared most fungal interactions with *G. breviflorus*, followed by *S. spruceanum* with *D. orobanchoides* and *V. aphylla*, and also *G. breviflorus* with *D. orobanchoides*. We observed that antagonistic plants as a group associate with fungi that are simultaneously linked to all mutualistic plants (Fig. **1b**), indicating that antagonistic plants are indirectly linked to any of the mutualistic plants detected in this study, which could therefore be the ultimate source of carbon for the antagonistic plants. Of all the possible connections between antagonistic and mutualistic plant species, *S. spruceanum* had the highest fungal overlap with *Clusia* sp., and then with *Tapirira* sp. and *Protium* sp.; *G. breviflorus* with *Aspidosperma* sp. and *Protium* sp., then with *Schadera* sp. and *Clusia* sp.; *D. orobanchoides* with *Tapirira* sp. and then with Fabaceae sp. 1; *V. parviflora* with *Aspidosperma* sp. and then with Araceae sp., Malpighiaceae sp., and *Clusia* sp; and *V. aphylla* with *Aspidosperma* sp. and then with *Paloue* sp.. These mutualistic plants are also those with highest number of fungal interactions overall (Fig. **3a**). Among the mutualistic plants, *Aspidosperma* sp. had the highest fungal overlap with *Protium* sp., Malpighiaceae sp. and Metteniusaceae sp., which are also among the highest ranked species in terms of number of associated fungi and phylogenetic diversity (Fig. **3**).

### Mutualistic and antagonistic plant-fungal interactions

We observed a significant correlation (Pearson correlation *r* = 0.63, *p* < 0.001, *t* = 8.27, *df* = 105) between the normalised degrees of the fungi in the antagonistic and mutualistic networks (Fig. **4**, Supporting Information Fig. **S3**). Fungi present in the mutualistic network but absent from the antagonistic network were generally characterized by a low normalized degree, except for two *Rhizophagus* taxa which were present in the majority of mutualistic plants (Fig. **4**). Moreover, we found a significant correlation of fungal ecological similarity between the mutualistic and antagonistic interactions (Mantel tests: Jaccard *r* = 0.20 ± 0.04, *p* = 0.002 ± 0.002; overlap *r* = 0.24 ± 0.04, *p* = 0.001), and also when accounting for the phylogenetic relatedness among the shared fungi (partial Mantel tests: Jaccard *r* = 0.18 ± 0.003, *p* = 0.018 ± 0.04; overlap *r* = 0.22 ± 0.04, *p* = 0.002 ± 0.003). We also observed that the fungi with the highest normalized degrees, both in the mutualistic and antagonistic network, are all members of the Glomeraceae (Fig. **4**).

**Figure 4:**
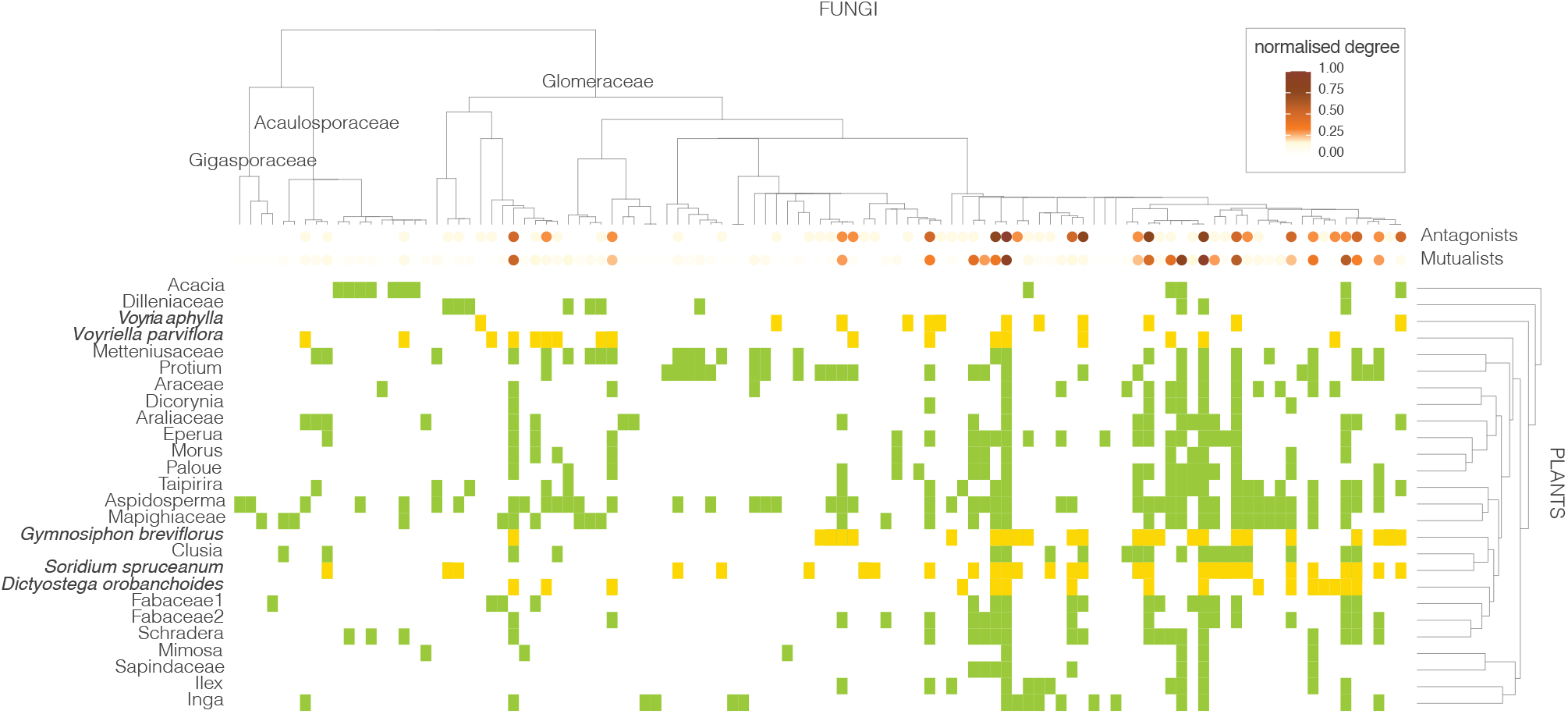
Plant-fungal interactions of mutualistic (green) and antagonistic (yellow) plants using a matrix rarefied to a depth of 844 reads. Phylogenetic relationships between the fungi are shown on top. Plant species are listed on the left; hierarchy clustered dendrogram based on the Bray-Curtis distance of their fungal communities is shown on the right. The intensity of the orange dots on the tips of the fungal phylogeny depicts the normalised degree (representing their interaction strength) of each fungus in each of the plant-fungal networks (i.e. in the antagonistic plants-fungi and mutualistic plants-fungi networks).

### Network analysis

The network analysis indicated that the presence of 2,560.39 ± 296.36 instances of the diamond-shaped module in the empirical networks across the rarefied matrices was overrepresented in relation to random expectations (*z-score* = 3.45 ± 0.40, *p* = 0.002 ± 0.003). The same analysis repeated for multiple rarefaction depths indicates that despite the number of diamond-shaped modules increases with increasing rarefaction depth, it does not impact the result that pairs of fungi share an antagonistic and a mutualistic plant more often than expected by chance (Supporting Information, Fig. **S4**).

## Discussion

We found that antagonistic plants as a group target approximately half of the fungi that are potentially available, yet this subset of fungi is associated with all mutualistic plants detected in this study. Thus, despite associating only with a subset of the local pool of fungi, the antagonists are still indirectly linked to every mutualistic plant. Fungi not detected in the roots of antagonists were generally connected to few mutualistic plants. Within the fungi that are shared between the mutualistic and antagonistic networks, we detected a significant ecological symmetry between the mutualistic and antagonistic interactions of the fungi: pairs of fungi that interact with similar sets of mutualistic plants also interact with overlapping antagonistic plants. The network analysis indicates that this pattern occurs more often than expected by chance. Based on the highest fungal overlap between each antagonistic species with subsets of highly connected mutualistic species, this pattern is mainly driven by an antagonist preference for fungi that are well-linked to specific mutualistic plants. These mutualistic plants are the ultimate sources of the carbon which antagonistic plants take up from the fungi shared between mutualists and antagonists. Therefore, we suggest that the observed pattern reflects a strategy in which the maintenance of antagonistic interactions is maximized by targeting well-linked mutualistic fungi, thereby minimizing the risk of carbon supply shortages.

### Fungal preferences of antagonistic plants

We found that plant species identity had a significant influence on the fungal community composition, regardless of the plant type (mutualist or antagonist), which indicates that these communities are non-random subsets of the local fungal taxon pool. This supports previous evidence that co-occurring plant species show differences in selectivity towards available arbuscular mycorrhizal fungi (Davison *et al*., 2011). Antagonistic plant species in particular, are known to select particular groups of fungi, often a narrower range than surrounding mutualistic plants (Bidartondo et al. 2002; Gomes et al. 2017). Here we observed that five co-occurring antagonistic plant species collectively associate with about half of the available fungal taxa (Fig. **4**). Antagonistic interactions can thus be supported by a relatively wide array of arbuscular mycorrhizal taxa, as shown previously (Merckx et al. 2012, Gomes *et al*., 2017b; Sheldrake *et al*., 2017). Although fungi from three different fungal families were detected in the roots of antagonistic plants, a clear preference for Glomeraceae taxa, and *Rhizophagus irregularis* relatives in particular, was observed. The taxa of this clade were the most frequently encountered in the roots of the autotrophic plants as well (Fig. **4**). *Rhizophagus* contains some of the most globally widespread and common arbuscular mycorrhizal fungi (Kivlin *et al*., 2011; Moora *et al*., 2011; Davison *et al*., 2015; Gomes *et al*., 2018) although this is mostly derived from studies in temperate areas. Our results indicate that the tropical rainforest offers no exception to this pattern. Glomeraceae are usually not only the most dominant clade in natural arbuscular mycorrhizal communities, often accounting for c. 70% of all species (Montesinos-Navarro *et al*., 2012), but they also have consistently been found to include the most generalist arbuscular mycorrhizal fungi in other network studies (e.g. Montesinos-Navarro *et al*., 2012; Chagnon *et al*., 2015; Chen *et al*., 2017). The ability to interact with many mutualistic plant species can be a potential reason for why antagonists generally target Glomeraceae fungi (Merckx *et al*., 2012; Renny *et al*., 2017). Ecological theory predicts that generalist species tend to have large distribution ranges (Brown, 1984) and, consequently, are less vulnerable to (local) extinction than specialized species (Schleuning *et al*., 2016). Therefore, associations to generalist fungi may be advantageous for the evolutionary persistence of antagonists. Furthermore, associations with multiple mutualistic plant partners may increase fungal resilience to disturbance, while mediating temporal fluctuations in carbon flow and interaction dynamics (Bennett *et al*., 2013) guaranteeing continuous carbon supply to the entire network without pronounced negative effects even in the presence of antagonists. In addition, in the context of mycorrhizal fungi, which can be linked to different plant species simultaneously (Montesinos-Navarro *et al*., 2012), generalist fungi are therefore likely to be more reliable carbon sources for antagonists. An alternative and perhaps not mutually exclusive explanation for why antagonistic plants preferentially target well-connected fungi may be that these fungi are less effective in detecting and excluding non-mutualistic plant partners (Bruns *et al*., 2002; Egger & Hibbett, 2004; Bidartondo, 2005; Walder & Van Der Heijden, 2015).

### Fungal links between mutualistic and antagonistic networks

We detected that pairs of fungi that interact with similar sets of mutualistic plants share links with overlapping antagonistic plants. Thus, there is a high level of interaction symmetry between mutualistic and antagonistic mycorrhizal networks. Also, we measured a significant influence of the fungal phylogenetic relationships on both the mutualistic and antagonistic interactions, showing that closely related fungi interact with similar mutualistic and antagonistic plants respectively. Because, biotic interactions are mediated by functional traits, and most functional traits are evolutionarily conserved, shared evolutionary history of fungi can serve as a proxy for functional similarity. We therefore hypothesize that both mutualistic and antagonistic interactions are shaped by evolutionary conserved functional traits of the fungi. In addition, the network analysis indicated that pairs of fungi share an antagonistic and a mutualistic plant more often than expected by chance. This analysis solely indicates that the diamond-shaped module is overrepresented in the empirical network (Fig. **2**), without reference to species degree nor to species identity. Yet, our results support that the observed pattern is driven by the tendency of antagonistic plants to target fungi that are well-linked to mutualistic plants (Fig. **1b**). The mutualistic plants with highest fungal overlap in relation to the antagonistic plants are among those with the highest ranked degree and phylogenetic species variability from the pool of detected mutualistic plant species (Fig. **3**). Moreover, fungi with the highest number of interactions in the mutualistic network are also among the best-connected fungi in the antagonistic network (Fig. **4**). Our findings thus reveal that antagonistic plants preferentially associate with fungi that are simultaneously linked to a wide range of mutualistic plants. Although many antagonistic plants share a large number of fungi with the mutualistic tree *Aspidosperma* sp., the mutualistic plants with highest normalised degree, each antagonistic plant species also indirectly associates with non-overlapping sets of mutualistic plants, as indicated by their divergent positions in the plant-plant interaction network (Fig. **2b**). This pattern suggests a potential strategy to avoid competition among antagonistic plant species while targeting well connected mutualistic plant species through shared fungi.

Our results indicate that antagonistic and mutualistic mycorrhizal networks are not linked randomly but are symmetrically influenced by mutualistic and antagonistic plant identity and fungal connectance. Whether these correlations are potentially influenced by spatial patterns in the distribution of mutualists, antagonists, and fungi remain to be determined. Our earlier work suggests that antagonistic mycorrhizal interactions can be better explained by understanding plant–plant interactions through sharing fungal associations (Gomes *et al*., 2017b), and indicates that antagonistic interactions may respond to an ecological mechanism driven by maximizing co-occurrence and avoiding competitive exclusion among antagonists. The results obtained here expand this view and suggest that not only plant-plant interactions drive patterns of co-occurrence of antagonistic plants, but also the indirect interactions between mutualistic and antagonistic plants through shared arbuscular mycorrhizal fungi.

### Potential sampling biases

Considering that rainforests are species-rich ecosystems (e.g. (Ter Steege *et al*., 2013)), it is possible that, despite our efforts, the sampling of the belowground diversity of arbuscular mycorrhizal fungi, and their plant partners remained incomplete. Furthermore, the representation of roots from antagonistic and mutualistic plants was necessarily imbalanced, as whole and partial root systems, respectively, were investigated. This made the use of read abundances to estimate interaction strengths impossible. While other studies have addressed the importance of sampling intensity, (i.e. the number of possible interactions per node which directly impacts the normalised degree per species), and network size (i.e. number of nodes involved) (Blüthgen *et al*., 2007; Dormann *et al*., 2009) on network metrics, the effect of species relationships is often elusive. In our approach, we accounted for the phylogenetic relationships between the fungi, as other studies have accounted for taxonomic diversity (Morris *et al*., 2014), and also we verified the consistency of our results across multiple rarefactions depths, by the use of multiple rarefied matrices at each depth (Gotelli & Colwell, 2001). This allows us to separate the statistical patterns in our data from the influence of sampling effort, both in terms of the plant species detected in the sampled roots, and in their potentially incomplete number of fungal associations. Furthermore, the number of interactions for mutualistic plants is moderately correlated with the phylogenetic diversity of the arbuscular mycorrhizal fungi associated with these tropical trees, while for antagonistic plants this correlation is not found. This further suggests that bias due to sampling effort is limited. It also indicates that antagonistic plants have specialised interactions, occupying particular fungal niches, which could have consequence for their occurrence and coexistence (Gomes *et al*., 2017b).

### Conclusions and future perspectives

To our knowledge, our study is the first to assess how antagonistic plant species are embedded in mutualistic mycorrhizal networks. We find that antagonistic plants as a group interact with half of the available fungal partners, and generally target fungi that are well-connected to mutualistic plants. Although antagonistic species show overlap in their fungal associations, we found that they are indirectly linked to different sets of mutualistic plants suggesting a potential mechanism to avoid competition by preferentially relying on different carbon sources (Gomes *et al*., 2017b). The phylogenetic relationships between the fungi, likely a proxy for fungal traits, have a significant influence on these non-random tripartite interactions. Therefore, we conclude that the persistence of antagonists in arbuscular mycorrhizal networks is dependent on particular well-connected ‘keystone’ mycorrhizal fungi, which provide the antagonists with carbon from a wide range of plants. Our observation that the way fungi connect mutualistic and antagonistic networks is not random and that well-connect fungal nodes in arbuscular mycorrhizal networks are more prone to be targeted by antagonists are similar to those of (Sauve *et al*., 2016) for a plant-pollinator-herbivore network when considering binary interactions. But further research is needed to assess whether this is a general feature of diffuse interactions, also when taking interaction strength into account. Our study emphasises the raising awareness of considering multiple interaction types simultaneously (e.g. antagonistic and mutualistic) to deepen our understanding of complex biodiversity patterns (Losapio *et al*., 2021).

In ectomycorrhizal systems mutualistic and achlorophyllous antagonistic plant species represent opposite extremes of a continuum of interaction outcomes and several plant species are able to combine photosynthesis and carbon uptake from fungi, a strategy known as ‘partial mycoheterotrophy’ (Selosse & Roy, 2009). Recent discoveries now indicate that partial mycoheterotrophy may be common among arbuscular mycorrhizal plants as well, and potentially many photosynthetic understory plants are able to take up carbon from associated arbuscular mycorrhizal fungi (Giesemann *et al*., 2021). It remains to be tested whether carbon uptake by these partially mycoheterotrophic plants occurs through sets of fungi, as suggested for fully mycoheterotrophic plant species.

## Acknowledgements

This research was supported by a Veni grant from the Netherlands Organisation of Scientific Research (NWO) to VSFTM (863.11.018) and field work was funded by the Royal Academy of Arts and Sciences of the Netherlands (KNAW) Ecology fund, grant J1606/Eco/G437 to SIFG. JB is supported by the Swiss National Science Foundation (Grant 31003A-169671 3). The authors thank the field assistance of Mélanie Roy, Vincent, Mathieu Gerard, the molecular lab assistance of Elza Duijm, and Hans ter Steege for verification of plant species occurrence in French Guiana. The authors also thank the valuable comments of four anonymous reviewers and editor Marc-André Selosse, which greatly improved the quality of the manuscript.

## Author contributions

SG, MF, JB, and VM designed the study and collected the materials. SG performed the laboratory work and data analysis. All authors contributed to the data analysis and the writing of the manuscript.

## Notes

### Competing Interest Statement

The authors have declared no competing interest.

