## Supplementary Information for "Antagonistic plants preferentially target arbuscular mycorrhizal fungi that are highly connected to mutualistic plants"

### *New Phytologist* Supporting Information

**Article title:** Mycoheterotrophic plants preferentially target arbuscular mycorrhizal fungi that are highly connected to autotrophic plants

**Authors:** Sofia IF Gomes, Miguel A Fortuna, Jordi Bascompte, Vincent SFT Merckx

**Article acceptance date:** Click here to enter a date.

The following Supporting Information is available for this article:

**Methods S1**| Effect of plant identity, plant type and subplot on the structure of fungal communities at the plant individual level.

**Methods S2** | Plant root identification.

**Fig S1** | Venn diagrams representing variation explained by plant species identity, plant type and subplot.

**Fig S2** | Phylogenetic signal analysis repeated on multiple rarefaction depths.

**Fig S3** | Motif analysis repeated on multiple rarefaction depths.

**Table S1** | Plant identity of autotrophic and mycoheterotrophic plants from this study.

**Methods S1** | Effect of plant identity, plant type and subplot on the structure of fungal communities at the plant individual level:

We explored the effects of plant identity, plant type (antagonist vs mutualist) and subplot origin on the fungal community composition obtained from the roots, at the individual plant level. For that, we rarefied each individual sample to 100 reads, and calculated the variance explained by these three factors using the function *varpart* using the *vegan* (Oksanen *et al.*, 2015) R package. We used two distance matrices by calculating the Bray-Curtis dissimilarities on the Hellinger transformed rarefied counts, and the Unifrac distance (Lozupone & Knight, 2005) to account for fungal relatedness. The variation partitioning analysis revealed that plant species identity captured most of the variation in the data, including the distinction between plant types (antagonist and mutualist), with a minor proportion of the variance explained by subplot (see Supporting Information, Figure S1), which can be expected due to the high heterogeneity of microbial communities in the soil at the local scale (Jacquemyn *et al.*, 2014). Then, to test for significance of association, we built a distance-based redundancy analysis (db-RDA) model for each matrix with these three factors, and performed model selection with the *ordi2step* function. The best model based showed that fungal communities were significantly structured by plant species identity, both considering their shared interactions only (*F* = 1.848, *P* = 0.001), or simultaneously accounting for the phylogenetic relationships between the shared fungi (*F* = 3.015, *P* = 0.001), explaining 44.8% or 57.7% of the variance in the data, respectively. The effect of plant type and plot were not significant in the db-RDA model.

**Methods S2** | Plant root identification:

Plant roots were identified to genus or family level based on *matK* or *trnL* sequencing using BLAST. The genus of the BLAST hit with the highest percent identity to the query sequence was used as the genus identification, unless this genus is not known to occur in the region. Query sequences with a similarity of 99% or higher were considered to belong to the same plant species. A time tree was constructed based on genus identification using TimeTree.org. For groups sequences that could not be identified to genus level with certainty, the genus of the best BLAST hit was used.

**Fig S1** | Venn diagrams representing variation explained by plant species identity, plant type and subplot using variation partitioning.

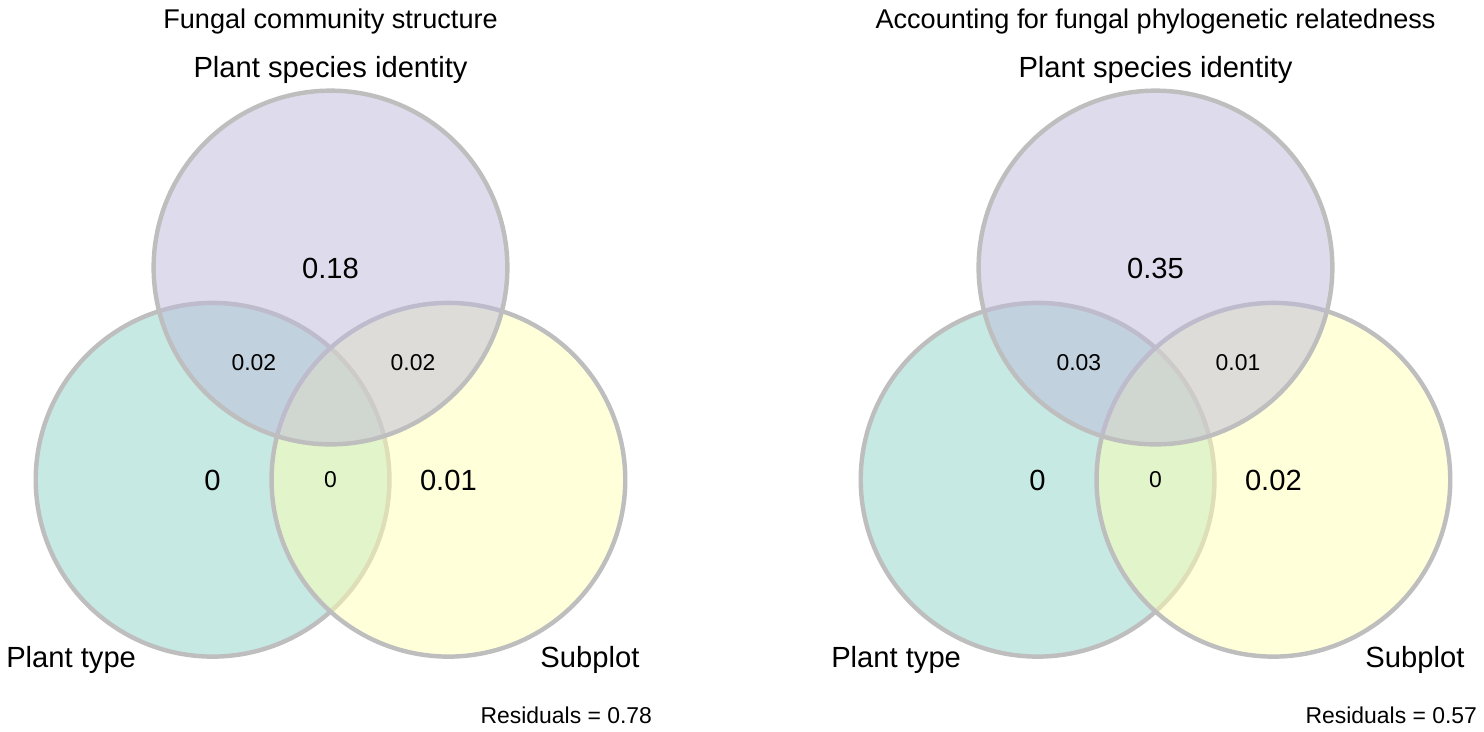

**Fig S2** | Phylogenetic signal analysis repeated on multiple rarefaction depths. Mean observed phylogenetic signal of fungi in the antagonistic (top) and mutualistic (bottom) networks for the 100 rarefaction matrices tested per rarefaction depth (left), and corresponding mean p-values (right).

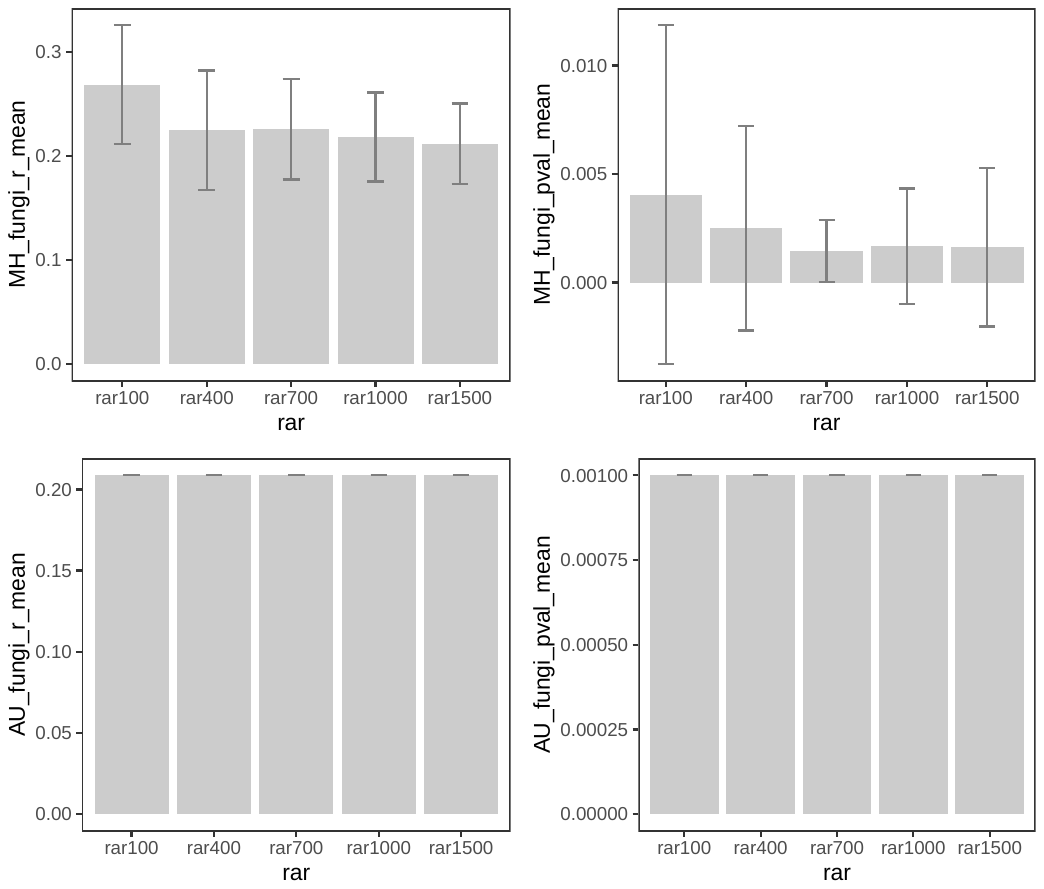

**Fig S3** | Relationship between fungal normalised degree in both the antagonistic and mutualistic networks (Pearson correlation r = 0.63, p < 0.001, t = 8.27, df = 105).

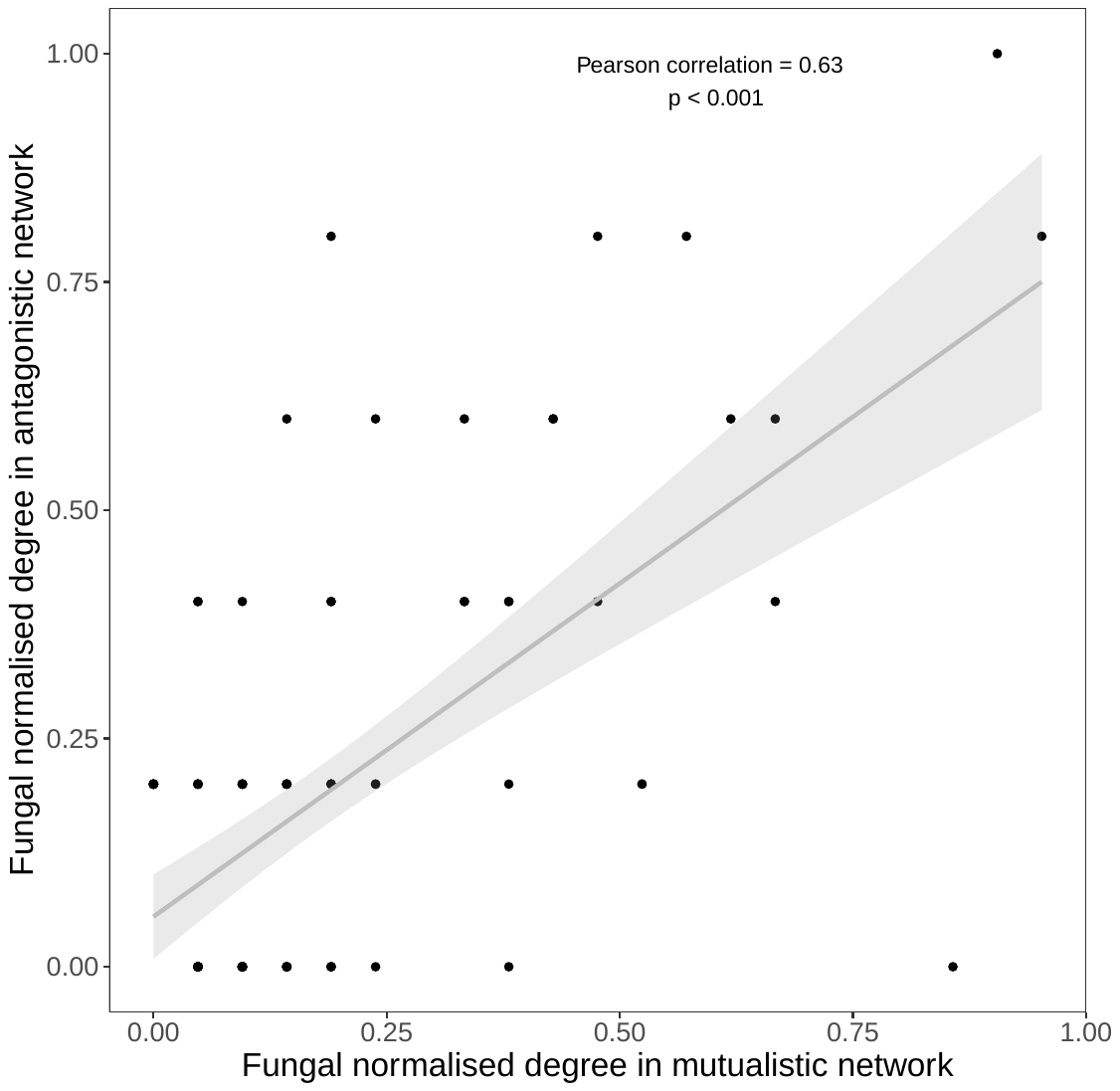

**Fig S4** | Motif analysis repeated on multiple rarefaction depths. Mean observed motifs for the 100 rarefaction matrices tested per rarefaction depth (left), and corresponding mean z-scores (right). Dashed lines represent the critical z-score values for 95% (y = 1.96) and 99% (y = 2.58) confidence levels.

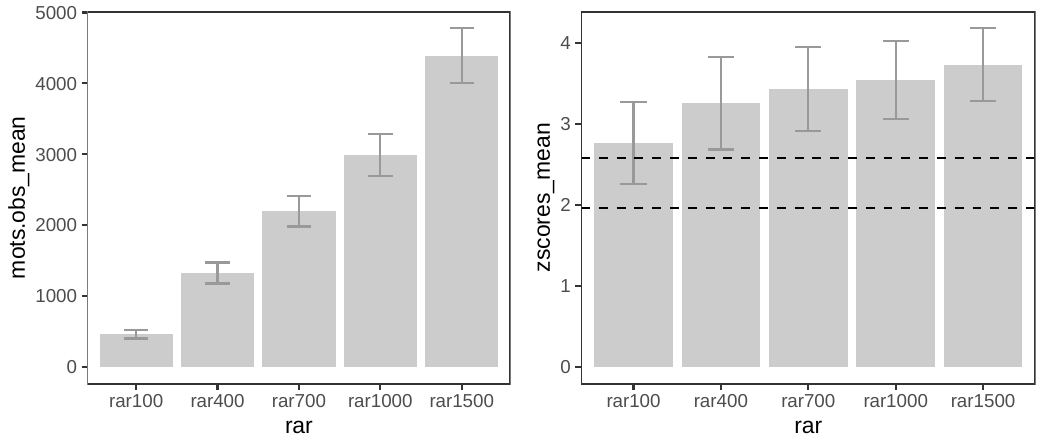

**Table S1** | Plant identity of autotrophic and mycoheterotrophic plants from this study. In total, we successfully extracted DNA from 123 autotrophic and 60 mycoheterotrophic individual plants. Of these, 99 individuals among 28 autotrophic, and 45 individuals among the five mycoheterotrophic plant species had Glomeromycotina reads. After removing samples with less than 500 reads, and only considering an OTU present in a sample when represented by at least 5 reads, we obtained a total of 82 individuals among 21 autotrophic, and 27 individuals among the five cheater plants. Rows in grey are excluded taxa due to the absence of Glomeromycotina reads; rows in orange are taxa excluded from subsequent analyses because had < 500 reads (total number of reads in autotrophic plants in the table excludes the taxa in orange).

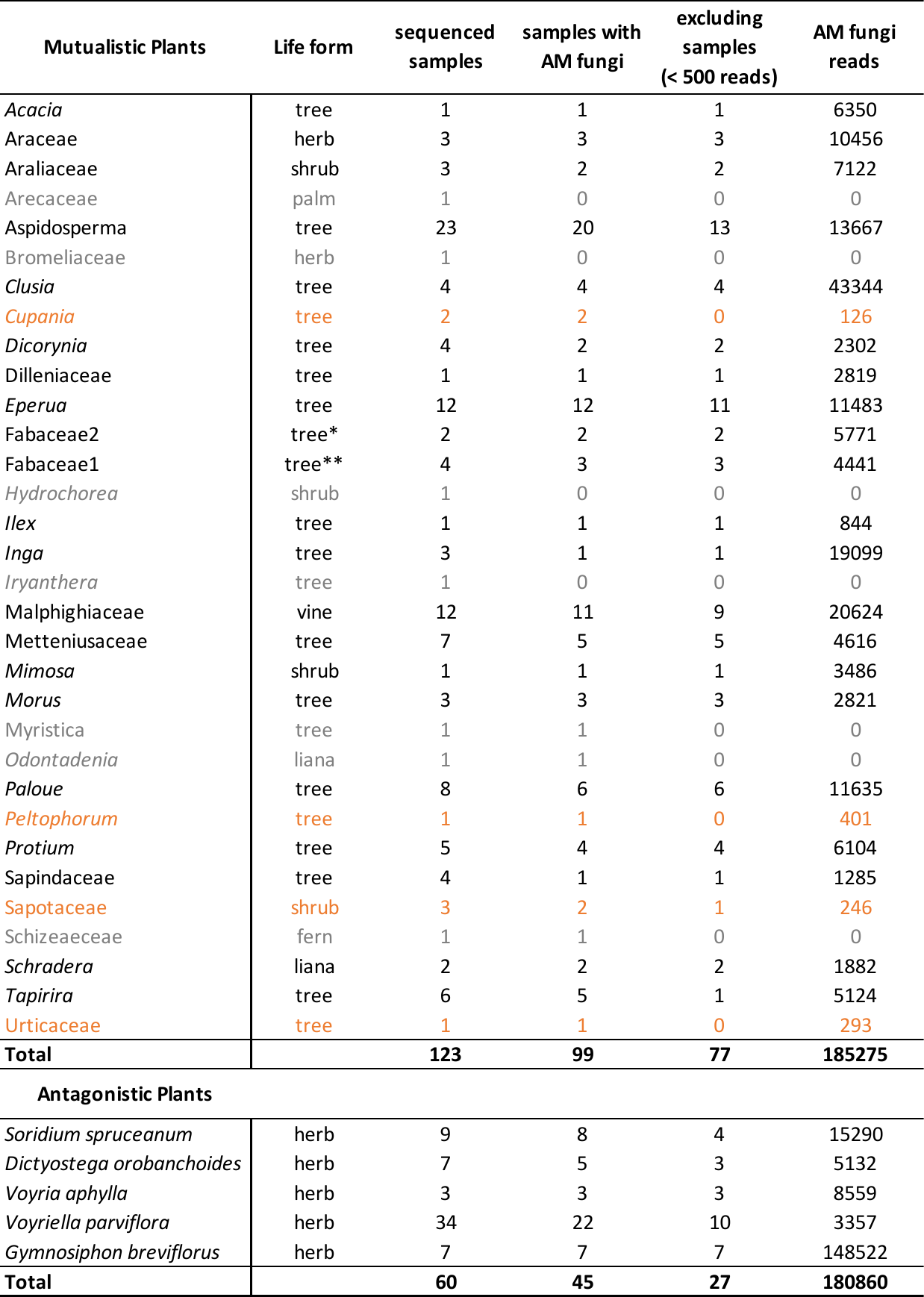

Identification of life form of these plants is based on the life form of their best Blast hit:

*best Blast hit (95.9%) is Martiodendron mediterraneum (does not occur in the region)

** best Blast hit (99.6%) is Acaciella chamalensis / Enterolobium cyclocarpum (does not occur in the region)
